# The pattern of Comorbidities and Multimorbidity among Colorectal Cancer Patients in Spain: CoMCoR study

**DOI:** 10.1101/526673

**Authors:** Miguel Angel Luque-Fernandez, Daniel Redondo-Sanchez, Miguel Rodríguez-Barranco, Maria Carmen Carmona-García, Rafael Marcos-Gragera, María-José Sánchez

## Abstract

Colorectal cancer is the most frequently diagnosed cancer in Spain. Cancer treatment and outcomes can be influenced by tumor characteristics, patient general health status and comorbidities. Numerous studies have analyzed the influence of comorbidity on cancer outcomes, but limited information is available regarding the frequency and distribution of comorbidities in colorectal cancer patients, particularly elderly ones, in the Spanish population. We developed a population-based study of all incident colorectal cancer cases diagnosed in Spain in 2011 to describe the frequency and distribution of comorbidities, as well as tumor and healthcare factors. Data were obtained from two population-based cancer registries and the complete revision of patients’ digitalized clinical records history. We then characterized the most prevalent comorbidities by patient, tumor and health care factors, as well as dementia and multimorbidity, and developed an interactive web application to visualize our findings. The most common comorbidities were diabetes (23.6%), chronic obstructive pulmonary disease (17.2%), and congestive heart failure (14.5%). Dementia was the most common comorbidity among patients aged ≥75 years. Patients with dementia had a 30% higher prevalence of being diagnosed at stage IV and the highest prevalence of emergency hospital admission after colorectal cancer diagnosis (33%). Colorectal cancer patients with dementia were nearly three times more likely to do not receive surgical treatment. Age ≥75 years, obesity, male sex, being a current smoker, having surgery more than 60 days after cancer diagnosis, and not receiving surgical treatment were associated with a higher prevalence of multimorbidity. Patients with multimorbidity aged ≥75 years showed a higher prevalence of hospital emergency admission followed by surgery the same day of the admission (37%). We found a consistent pattern in the distribution and frequency of comorbidities and multimorbidity among colorectal cancer patients. The high frequency of stage IV diagnosis among patients with dementia and the high proportion of older patients not receiving surgical treatment are significant findings that require policy actions.

## Introduction

Cancer accounted for 9.6 million deaths globally in 2018, and was the second most common cause of death in the world [1]. Colorectal cancer (CRC) is the most frequently diagnosed cancer in Spain, with 44,937 estimated new CRC cases in 2019 [2, 3]. Despite the high prevalence of CRC in the elderly, the inclusion of this cohort in clinical trials is disproportionately low [4]. In addition to clinical and pathological characteristics of the tumor, general health status and comorbidities of patients also influence cancer treatment and outcomes. Comorbidity describes the existence of a long-term health condition or disorder in the presence of a primary disease of interest, such as cancer [5], whereas multimorbidity refers to the existence of more than one comorbid condition [6]. Comorbidity and multimorbidity are increasingly seen as a problem of the elderly, but have also been reported as occurring more often and at a younger age in patients of lower socioeconomic status [7, 8]. The presence of comorbidities can influence treatment options, and therefore should be thoroughly evaluated when studying prognosis, outcomes, and mortality in cancer patients. Despite the coexistence of health conditions being commonplace, the guidelines and delivery of care appear to be focused on single disease management [9, 10]. However, effective management of comorbid conditions is important in maintaining patients’ optimal health status, as the presence of one could contribute to the development of another [11], and decisions regarding cancer treatment require the consideration of patients’ comorbidities [12–14]. Furthermore, post-operative complications have been reported as higher in patients with comorbidity [15], and certain comorbid conditions have been linked to adverse outcomes following surgery for cancer [12, 16].

As noted above, there is consistent evidence on the influence of comorbidities on cancer outcomes, but little is known about them in CRC patients. Thus, we aimed to describe and characterize the frequency and distribution of comorbidities and multimorbidity by patient, tumor and health care factors among all CRC incident cases diagnosed in Granada and Girona (Spain) in 2011.

## Materials and methods

### Study design, participants, data, and setting

We conducted a population-based study including all the CRC incident cases diagnosed in 2011 in two Spanish provinces (Girona and Granada), registered with codes (C18-C21) according to the International Classification of Diseases for Oncology, 3^rd^ Edition, (ICD-O-3), and followed up until December 31, 2016.

Data were obtained retrospectively from two Spanish population-based cancer registries and the revision of the complete and digitalized patients’ clinical records history, including primary care, out, and inpatient hospital information. The data collection followed a detailed protocol from the European High Resolution studies collaboration (TRANSCAN-HIGHCARE project within ERA-Net) [17]. We recorded information regarding the cancer stage at diagnosis (TNM staging system, 7^th^ edition), cancer diagnostic exams, tumor morphology, cancer treatment, patients’ comorbidities, performance status, and vital status. All recorded comorbidities were extracted 6 months before the index cancer was diagnosed, based on a standardized protocol published elsewhere [18]. All information was classified as either patient, tumor, or healthcare factors.

The study proposal (CP17/00206 CoMCoR study) was approved by an internal review board from the Andalusian School of Public Health. The ethics review committee from the Department of Health of the Andalusian regional government approved the study with internal number 0072-N-18. No samples were used, all data were fully anonymized, and the informed consent was waived.

### Variables related to the patient’s characteristics

We recorded patient’s age, sex, smoking status, body mass index (BMI), performance status, comorbidities, and multimorbidity. Age at diagnosis was categorized into four age groups: <55, 55-64, 65-74, and ≥75 years. Smoking status was categorized as current, previous, and never smoker. BMI was categorized as underweight-normal (<25.0 kg/m^2^), overweight (≥25.0 kg/m^2^ and <30 kg/m^2^), and obese (≥30 kg/m^2^). We combined the underweight with normal weight category because of data sparsity (i.e., only 4 patients). Patients’ performance status was ascertained based on the retrospective revision of their clinical history and the Eastern Cooperative Oncology Group (ECOG) scale. ECOG was categorized as normal (0); restricted but able to carry out light work (1); restricted, unable to work but capable of self-care (2); restricted, capable of limited self-care (3); and disabled (4) [19]. Comorbidities were classified based on the Royal College of Surgeons modified Charlson score that reduces the number of comorbidities to 12 (myocardial infarction, congestive heart failure, peripheral vascular disease, cerebrovascular disease, dementia, chronic obstructive pulmonary disease (COPD), rheumatic disease, liver disease, diabetes mellitus, hemiplegia/paraplegia, renal disease and AIDS/HIV), removing some categories such as peptic ulcer disease (since it is not considered a chronic disease anymore), and grouping diseases together (e.g., diabetes mellitus codes with or without complications are grouped into one category). Furthermore, the score does not assign weights to comorbidities, and instead categorizes the number of comorbidities in three different groups: 0, 1, and ≥2 as a multimorbidity indicator [20].

### Variables related to tumor characteristics

We recorded the tumor topography, morphology, grade of differentiation, and stage at diagnosis. The final stage variable was defined as the combination of clinical and pathological TNM stages and categorized into five groups, based on the 7^th^ edition of the TNM manual. Topography, grade of differentiation, and morphology were coded according to ICD-O-3.

### Variables related to healthcare provision factors

We recorded the type of hospital admission, surgery, type of surgery, and time to surgery. Type of hospital admission indicated whether cancer patients had an emergency or planned admission. The type of surgery was dichotomized as major or minor was assessed based on the Classification of Interventions and Procedures (fourth version, ‘OPCS-4’). Major surgery included: anterior resection (H333-34, H336, H3380); Hartman (H335); abdomino-perineal resection (H331); right hemicolectomy (H061-64, H068-H079); left hemicolectomy (H091-99); segmental resection (H048, H081-89, H101-109, H111-118, H339); and total colectomy (H051-59). Endoscopic resection, endoscopic polypectomy and transanal resection were classified as minor surgery. The time to surgery was noted as the number of days from the date of cancer diagnosis to the date patients had the surgical intervention with curative or palliative intent. Time to surgery was categorized into five groups (0, 1 to <14, 14 to 30, 31 to 59 and ≥60 days). Emergency surgery was defined as surgery offered on the same day of an emergency hospital admission.

### Statistical analysis

First, we calculated the prevalence of each of the 12 different comorbidities for the all CRC patients. However, we only presented the results for 10 comorbidities as HIV and hemiplegia/paraplegia were only represented for 4 and 3 cases, respectively. Then, we calculated the frequency and distribution of comorbidities by patient, tumor and healthcare factors using counts and proportions. The Chi-square, Fisher’s exact, and score tests were used for statistical hypothesis testing. We assumed missing data, in a completely at random pattern, and thus performed a complete case analysis. Afterward, we computed unadjusted, sex-adjusted, and age-adjusted comorbidity prevalence ratios (PRs) with 95% confidence intervals (CIs) by patient, tumor, and healthcare factors. Generalized linear models with Poisson distribution and log link were fitted for the five most common comorbidities plus dementia. To estimate the comorbidities PR we included the specific comorbidity indicator as the dependent variable and the patient, tumor, and health care factors as the independent variables [21]. To describe the prevalence of multimorbidity (≥2 chronic conditions vs. non-comorbidities) by patients, tumor and health care factors, we used a multinomial logistic regression using the Royal College of Surgeons modified Charlson score as the dependent variable, with patient, tumor, and health care factors as independent variables. Then, we derived unadjusted, age-adjusted, and sex-adjusted PRs with 95% CIs. Finally, we developed an open source web application using advanced visualization tools (radar plots, heat maps and forest plots) [22] to present the complete results of the study, available at http://watzilei.com/shiny/CoMCoR/. Furthermore, we created a GitHub repository where the code used to develop the analysis and the web application can be accessed for reproducibility (https://github.com/migariane/CoMCoR). We used Stata v.15.1 (StataCorp, College Station, Texas, U.S.) and R v.3.5 (R Foundation for Statistical Computing, Vienna, Austria) for statistical analysis.

## Results

Table 1 shows the prevalence of comorbidities among CRC patients at least 6 months before the cancer diagnosis, ordered by frequency. Diabetes mellitus, COPD, and congestive heart failure were the most common comorbidities among CRC patients (24%, 17%, and 15%, respectively) (Table 1).

**Table 1.** Prevalence of comorbidities ordered by frequency among all incident colorectal cancer patients in Granada and Girona, 2011, n = 1,061

|  | Comorbidities Prevalence |  |  |  |  |  |  |  |  |  |  |  |
| --- | --- | --- | --- | --- | --- | --- | --- | --- | --- | --- | --- | --- |
|  | IX | VI | II | III | VII | XI | I | IV | VIII | V | XII | X |
|  | n(%) | n(%) | n(%) | n(%) | n(%) | n(%) | n(%) | n(%) | n(%) | n(%) | n(%) | n(%) |
| Yes | 250(23.6) | 182(17.2) | 154(14.5) | 124(11.7) | 104(9.8) | 92(8.7) | 67(6.3) | 65(6.1) | 56(5.3) | 48(4.5) | 4(0.4) | 3(0.3) |
| No | 790(74.5) | 859(81.0) | 887(83.6) | 917(86.4) | 937(88.3) | 947(89.3) | 974(91.8) | 976(92.0) | 985(92.8) | 992(93.5) | 1036(97.6) | 1037(97.7) |
| Unknown | 21(2.0) | 20(1.9) | 20(1.9) | 20(1.9) | 20(1.9) | 22(2.1) | 20(1.9) | 20(1.9) | 20(1.9) | 21(2.0) | 21(2.0) | 21(2.0) |

**Comorbidities:** **I** Myocardial infarction; **II** Congestive heart failure; **III** Peripheral vascular disease; **IV** Cerebrovascular disease; **V** Dementia; **VI** Chronic obstructive pulmonary disease; **VII** Rheumatic disease; **VIII** Liver disease; **IX** Diabetes mellitus; **X** Hemiplegia/paraplegia; **XI** Renal disease; **XII** AIDS/HIV.

### Patient and tumor characteristics

Table 2 shows the distribution of patient, tumor, and healthcare characteristics for all the colorectal cancer patients under study (n=1.061). The degree of completeness of patients’ medical records was high. It was reflected by a relatively low percentage of missing information for important variables such as cancer stage (5.6%), comorbidities (2.2%), and surgery (0.7%). More than half (59%) of colorectal cancer patients had one or more comorbidities 6 months before cancer diagnosis, and 30% had multimorbidity (i.e. CRC plus two or more comorbidities). The maximum number of comorbidities we found was six (only four CRC patients). Men represented 61% of the cohort, 67% of patients were age >65 years, 12% had a restricted performance status, slightly more than half of them were previous or current smokers (52%), and 49% were overweight or obese. The prevalence of the different tumor locations was 34% in the right colon, 32% in the left colon, and 33% in the rectum. The differentiation of the tumor was mostly grade two (56%); however, 19% of the tumors were not graded. Only 16% of colorectal cancer patients had a stage I tumor at diagnosis, while more than 50% of the cases were identified as stage III/IV. Six percent of patients had missing stage information. The type of hospital admission was principally planned (65%), and almost one out of five patients were admitted after visiting the hospital emergency department. Surgery was performed in 83% of the patients, and the most frequent type of surgery was major surgery (77%). The time to surgery exceeded 60 days for 26% of the patients. Sixteen percent of the colorectal cancer cases had emergency surgery (Table 2).

**Table 2.** Distribution of patient, tumor and healthcare characteristics among all incident colorectal cancer patients in Granada and Girona, 2011, n = 1,061

| Patient's characteristics |  | N(%) | Healthcare factors |  | N(%) | Tumor's characteristics |  | N(%) | Comorbidities* |  | N(%) |
| --- | --- | --- | --- | --- | --- | --- | --- | --- | --- | --- | --- |
| Age in years |  |  | Type hospital admission |  |  | Anatomical subsite |  |  | Multimorbidity Prevalence |  |  |
| <55 |  | 130(12.3) | Emergency |  | 183(17.2) | Right colon |  | 357(33.6) | None |  | 413(38.9) |
| 55 - 64 |  | 219(20.6) | Planned |  | 693(65.3) | Left colon |  | 340(32.1) | One |  | 301(28.4) |
| 65 - 74 |  | 272(25.6) | Missing |  | 185(17.4) | Colon Unspecified |  | 11(1.0) | Two |  | 190(17.9) |
| ≥75 |  | 440(41.5) | **Surgery** |  |  | Rectal |  | 353(33.3) | Three |  | 89(8.4) |
| **Sex** |  |  | Yes |  | 879(82.8) | **Grade of differentiation** |  |  | Four |  | 30(2.8) |
| Male |  | 644(60.7) | No |  | 175(16.5) | One |  | 168(15.8) | Five |  | 11(1.0) |
| Female |  | 417(39.3) | Missing |  | 7(0.7) | Two |  | 596(56.2) | Six |  | 4(0.4) |
| **Performance status ECOG score** |  |  | **Type of Surgery** |  |  | Three |  | 90(8.5) | Missing |  | 23(2.2) |
| Normal (0) |  | 259(24.4) | Not done |  | 175(16.5) | Four |  | 7(0.6) |  |  |  |
| Restricted but able to carry out light work (1) |  | 423(39.9) | Major |  | 816(76.9) | Missing |  | 200(18.9) |  |  |  |
| Restricted, unable to work but capable of selfcare (2) |  | 83(7.8) | Minor |  | 43(4.1) | **Stage TNM** |  |  |  |  |  |
| Restricted, capable of limited selfcare (3) |  | 35(3.3) | Done but unknown type |  | 20(1.9) | I |  | 168(15.8) |  |  |  |
| Disabled (4) |  | 6(0.6) | Missing |  | 7(0.7) | II |  | 281(26.5) |  |  |  |
| Missing |  | 255(24.0) | **Time to surgery in months** |  |  | III |  | 285(26.9) |  |  |  |
| **Smoking status** |  |  | Emergency 0 days |  | 171 (16.1) | IV |  | 267(25.2) |  |  |  |
| Current |  | 130(12.3) | 1 to <14 days |  | 115(10.8) | Missing |  | 60(5.6) |  |  |  |
| Previous |  | 298(28.1) | 14 to 30 days |  | 124(11.7) |  |  |  |  |  |  |
| Never |  | 505(47.6) | 31 to 59 days |  | 188(17.7) |  |  |  |  |  |  |
| Missing |  | 127(12.0) | 60 and more days |  | 280(26.4) |  |  |  |  |  |  |
| **BMI in kg/m2** |  |  | Missing |  | 8 (0.8) |  |  |  |  |  |  |
| <25 |  | 226(21.3) | No surgery |  | 175 (16.5) |  |  |  |  |  |  |
| 25.0 - 29.9 |  | 327(30.8) |  |  |  |  |  |  |  |  |  |
| ≥30 |  | 193(18.2) |  |  |  |  |  |  |  |  |  |
| Missing |  | 315(29.7) |  |  |  |  |  |  |  |  |  |

\* Comorbidity score based on: Armitage JN, van der Meulen JH. Identifying co-morbidity in surgical patients using administrative data with the Royal College of Surgeons Charlson Score. The British journal of surgery. 2010 May;97(5):772-81.

Figure 1 contrasts the distribution of the pattern in the distribution of the 10 most common comorbidities by sex. The most common condition among men with comorbidities was COPD, represented by the highest frequency in the plot. The frequency of COPD among men was 80% versus 20% in women. Among women with comorbidities the most common conditions were rheumatologic disease and dementia with 60% and 56% versus 40% and 43% in men, respectively.

**Fig 1.**
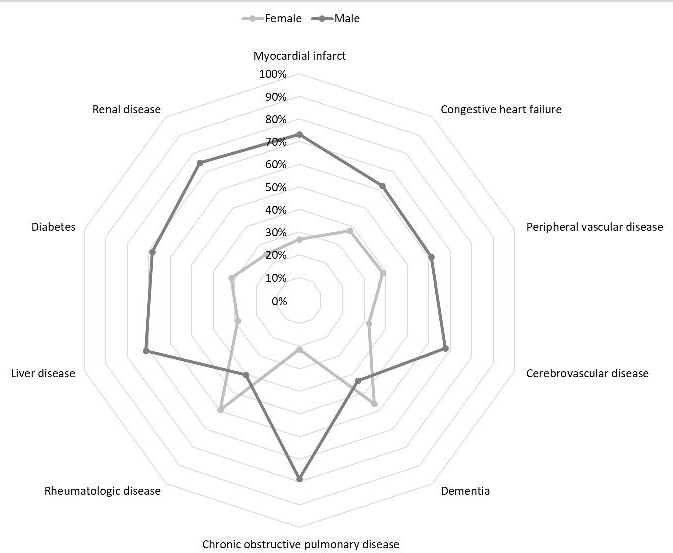
Radar plot displaying the prevalence of comorbidities by sex among all incident colorectal cancer patients in Granada and Girona, 2011, n = 1,061.

Figure 2 shows the distribution of the prevalence of the top-ten comorbidities by age. The most common comorbidity among elderly (age ≥75 years) was dementia and liver disease among patients aged <55 years.

**Fig 2.**
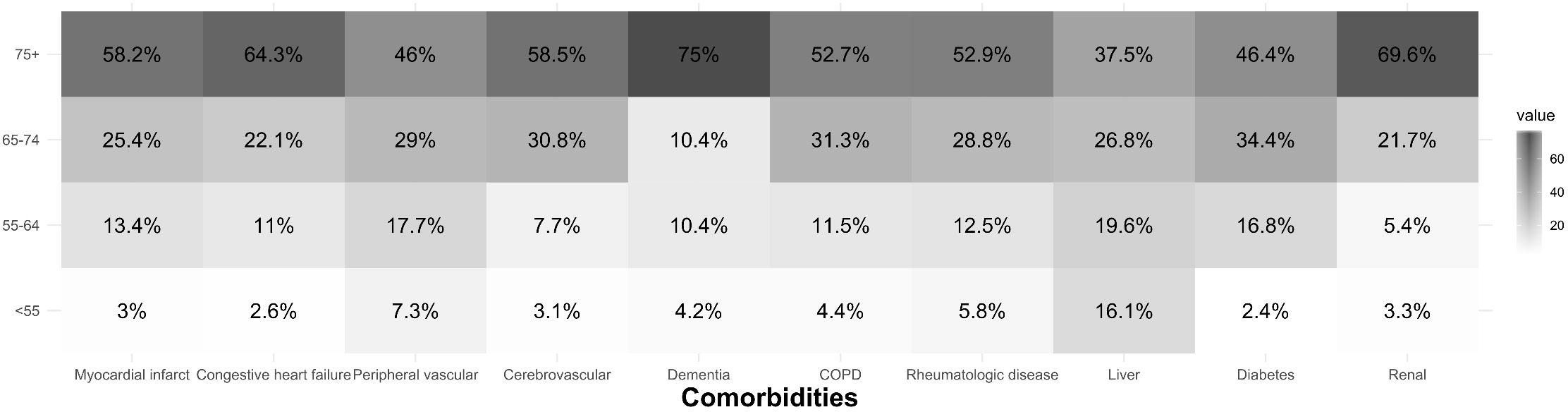
Heat map displaying the prevalence of comorbidities by age among all incident colorectal cancer patients in Granada and Girona, 2011, n = 1,061. Comorbidities: I Myocardial infarct; II Congestive heart failure; III Peripheral vascular disease; IV Cerebrovascular disease; V Dementia; VI Chronic pulmonary disease; VII Rheumatic disease; VIII Liver disease; IX Diabetes mellitus; XI Renal disease.

Table 3 shows the frequency and crude prevalence ratio of comorbidities for the five most common comorbidities plus dementia by tumor, patient, and health care factors. Supporting information Table S1 shows sex-adjusted and age-adjusted comorbidity prevalence ratios by tumor, patient, and health care factors. The complete distribution of comorbidities is provided as supporting information (Supplementary Tables S2, S3, and S4).

**Table 3.** Distribution and frequency of the top five comorbidities plus dementia and associated prevalence ratios by patient, tumor and healthcare characteristics among all incident colorectal cancer patients in Granada and Girona, 2011, n = 1,061.

| Patient's factors | Total | II |  |  | III |  |  | V |  |  | VI |  |  | VII |  |  | IX |  |  |
| --- | --- | --- | --- | --- | --- | --- | --- | --- | --- | --- | --- | --- | --- | --- | --- | --- | --- | --- | --- |
|  |  | n(%) | P(%) | PR(95%CI) | n(%) | P(%) | PR(95%CI) | n(%) | P(%) | PR(95%CI) | n(%) | P(%) | PR(95%CI) | n(%) | P(%) | PR(95%CI) | n(%) | P(%) | PR(95%CI) |
| **Age in years** |  |  |  |  |  |  |  |  |  |  |  |  |  |  |  |  |  |  |  |
| <55 | 130 | 4(2.6) | 3.1 | (Reference) | 9(7.3) | 6.9 | (Reference) | 2(4.2) | 1.5 | (Reference) | 8(4.4) | 6.2 | (Reference) | 6(5.8) | 4.6 | (Reference) | 6(2.4) | 4.6 | (Reference) |
| 55 - 64 | 219 | 17(11) | 7.8 | 2.5(0.9, 7.4) | 22(17.7) | 10.1 | 1.5(0.7, 3.1) | 5(10.4) | 2.3 | 1.5(0.3, 7.6) | 21(11.5) | 9.7 | 1.6(0.7, 3.4) | 13(12.5) | 6.0 | 1.3(0.5, 3.3) | 42(16.8) | 19.4 | 4.2(1.8, 9.6) |
| 65 - 74 | 272 | 34(22.1) | 12.6 | 4.1(1.5, 11.3) | 36(29.0) | 13.4 | 1.9(1.0, 3.9) | 5(10.4) | 1.9 | 1.2(0.2, 6.1) | 57(31.3) | 21.2 | 3.4(1.7, 7.0) | 30(28.8) | 11.2 | 2.4(1.0, 5.7) | 86(34.4) | 32.0 | 6.9(3.1, 15.4) |
| ≥75 | 440 | 99(64.3) | 23.3 | 7.6(2.8, 20.2) | 57(46.0) | 13.4 | 1.9(1.0, 3.8) | 36(75.0) | 8.5 | 5.5(1.3, 22.6) | 96(52.7) | 22.6 | 3.7(1.8, 7.3) | 55(52.9) | 12.9 | 2.8(1.2, 6.4) | 116(46.4) | 27.3 | 5.9(2.7, 13.1) |
| **Sex** |  |  |  |  |  |  |  |  |  |  |  |  |  |  |  |  |  |  |  |
| Male | 644 | 96(62.3) | 15.2 | (Reference) | 76(61.3) | 12.0 | (Reference) | 21(43.8) | 3.3 | (Reference) | 143(78.6) | 22.7 | (Reference) | 42(40.4) | 6.7 | (Reference) | 171(68.4) | 27.1 | (Reference) |
| Female | 417 | 58(37.7) | 14.1 | 0.9(0.7, 1.3) | 48(38.7) | 11.7 | 1.0(0.7, 1.4) | 27(56.3) | 6.6 | 2(1.1, 3.5) | 39(21.4) | 9.5 | 0.4(0.3, 0.6) | 62(59.6) | 15.1 | 2.3(1.6, 3.3) | 79(31.6) | 19.3 | 0.7(0.6, 0.9) |
| **Performance status** |  |  |  |  |  |  |  |  |  |  |  |  |  |  |  |  |  |  |  |
| Normal (0) | 259 | 20(16.0) | 7.8 | (Reference) | 12(12.6) | 4.7 | (Reference) | 1(3.3) | 0.4 | (Reference) | 25(17.9) | 9.7 | (Reference) | 19(19.0) | 7.4 | (Reference) | 45(22.3) | 17.5 | (Reference) |
| Restricted but able to carry out light work (1) | 423 | 68(54.4) | 16.1 | 2.1(1.3, 3.3) | 62(65.3) | 14.7 | 3.2(1.7, 5.7) | 13(43.3) | 3.1 | 7.9(1.0, 60.4) | 89(63.6) | 21.0 | 2.2(1.4, 3.3) | 66(66.0) | 15.6 | 2.1(1.3, 3.4) | 117(57.9) | 27.7 | 1.6(1.2, 2.1) |
| Restricted, unable to work but capable of selfcare (2) | 83 | 21(16.8) | 25.6 | 3.3(1.9, 5.8) | 10(10.5) | 12.2 | 2.6(1.2, 5.8) | 8(26.7) | 9.8 | 25.2(3.2, 198.3) | 16(11.4) | 19.5 | 2.0(1.1, 3.6) | 9(9.0) | 11.0 | 1.5(0.7, 3.2) | 25(12.4) | 30.5 | 1.7(1.1, 2.7) |
| Restricted, capable of limited selfcare (3) | 35 | 12(9.6) | 34.3 | 4.4(2.4, 8.2) | 9(9.5) | 25.7 | 5.5(2.5, 12.2) | 6(20.0) | 17.1 | 44.2(5.5, 356.6) | 8(5.7) | 22.9 | 2.4(1.2, 4.8) | 4(4.0) | 11.4 | 1.6(0.6, 4.3) | 14(6.9) | 40.0 | 2.3(1.4, 3.7) |
| Disabled (4) | 6 | 4(3.2) | 66.7 | 8.6(4.2, 17.4) | 2(2.1) | 33.3 | 7.2(2.0, 25.2) | 2(6.7) | 33.3 | 86.0(9.0, 82.4) | 2(1.4) | 33.3 | 3.4(1.0, 11.3) | 2(2.0) | 33.3 | 4.5(1.3, 15.2) | 1(0.5) | 16.7 | 1.0(0.2, 5.8) |
| **Smoking status** | 255 |  |  |  |  |  |  |  |  |  |  |  |  |  |  |  |  |  |  |
| Current | 130 | 12(8.9) | 9.2 | 0.7 (0.4, 1.3) | 15(13.0) | 11.5 | 0.9 (0.5, 1.6) | 5(12.8) | 3.8 | 0.8 (0.3, 2.0) | 35(20.7) | 26.9 | 2.5 (1.6, 3.9) | 9(9.8) | 6.9 | 0.5 (0.3, 1.0) | 31(14.0) | 23.8 | 1.2 (0.8, 1.7) |
| Previous | 298 | 55(40.7) | 18.5 | 1.4 (1.0, 2.0) | 35(30.4) | 11.7 | 0.9 (0.6, 1.4) | 9(23.1) | 3.0 | 0.6 (0.3, 1.3) | 80(47.3) | 26.8 | 2.5 (1.8, 3.5) | 16(17.4) | 5.4 | 0.4 (0.2, 0.7) | 87(39.2) | 29.2 | 1.4 (1.0, 1.9) |
| Never | 505 | 68(50.4) | 13.5 | (Reference) | 65(56.5) | 12.9 | (Reference) | 25(64.1) | 5.0 | (Reference) | 54(32.0) | 10.7 | (Reference) | 67(72.8) | 13.3 | (Reference) | 104(46.8) | 20.6 | (Reference) |
| **BMI in kg/m2** |  |  |  |  |  |  |  |  |  |  |  |  |  |  |  |  |  |  |  |
| <25 | 226 | 17(19.5) | 7.5 | (Reference) | 23(23.5) | 10.2 | (Reference) | 12(41.4) | 5.3 | (Reference) | 40(30.8) | 17.7 | (Reference) | 22(32.4) | 9.7 | (Reference) | 40(23) | 17.7 | (Reference) |
| 25.0 - 29.9 | 327 | 40(46.0) | 12.2 | 1.6(0.9, 2.8) | 42(42.9) | 12.8 | 1.3(0.8, 2.0) | 10(34.5) | 3.1 | 0.6(0.3, 1.3) | 41(31.5) | 12.5 | 0.7(0.5, 1.1) | 25(36.8) | 7.6 | 0.8(0.5, 1.4) | 74(42.5) | 22.7 | 1.3(0.9, 1.8) |
| ≥30 | 193 | 30(34.5) | 15.5 | 2.1(1.2, 3.6) | 33(33.7) | 17.1 | 1.7(1.0, 2.8) | 7(24.1) | 3.6 | 0.7(0.3, 1.7) | 49(37.7) | 25.4 | 1.4(1.0, 2.1) | 21(30.9) | 10.9 | 1.1(0.6, 2.0) | 60(34.5) | 31.1 | 1.8(1.2, 2.5) |

|  |  | II |  |  | III |  |  | V |  |  | VI |  |  | VII |  |  | IX |  |  |
| --- | --- | --- | --- | --- | --- | --- | --- | --- | --- | --- | --- | --- | --- | --- | --- | --- | --- | --- | --- |
|  |  | n(%) | P(%) | PR<br>(95%CI) | n(%) | P(%) | PR<br>(95%CI) | n(%) | P(%) | PR<br>(95%CI) | n(%) | P(%) | PR<br>(95%CI) | n(%) | P(%) | PR<br>(95%CI) | n(%) | P(%) | PR<br>(95%CI) |
| Tumor factors | Total |  |  |  |  |  |  |  |  |  |  |  |  |  |  |  |  |  |  |
| Anatomical Site |  |  |  |  |  |  |  |  |  |  |  |  |  |  |  |  |  |  |  |
| Right colon | 357 | 55(35.7) | 15.7 | (Reference) | 44(35.5) | 12.5 | (Reference) | 21(43.8) | 6.0 | (Reference) | 67(36.8) | 19.1 | (Reference) | 31(29.8) | 8.8 | (Reference) | 98(39.2) | 28.8 | (Reference) |
| Left colon | 340 | 50(32.5) | 14.9 | 1.0(0.7, 1.4) | 41(33.1) | 12.2 | 1.0(0.7, 1.5) | 11(22.9) | 3.3 | 0.5(0.3, 1.1) | 63(34.6) | 18.8 | 1.0(0.7, 1.3) | 33(31.7) | 9.9 | 1.1(0.7, 1.8) | 74(29.6) | 22.1 | 0.8(0.6, 1.0) |
| Colon Unspecified | 11 | 2(1.3) | 28.6 | 1.8(0.6, 6.0) | 1(0.8) | 14.3 | 1.1(0.2, 7.1) | 0(0) | - | - | 1(0.5) | 14.3 | 0.7(0.1, 4.7) | 1(1.0) | 14.3 | 1.6(0.3, 10.2) | 2(0.8) | 28.6 | 1.0(0.3, 3.3) |
| Rectal | 353 | 47(30.5) | 13.5 | 0.9(0.6, 1.2) | 38(30.6) | 10.9 | 0.9(0.6, 1.3) | 16(33.3) | 4.6 | 0.8(0.4, 1.4) | 51(28.0) | 14.7 | 0.8(0.6, 1.1) | 39(37.5) | 11.2 | 1.3(0.8, 2.0) | 76(30.4) | 21.8 | 0.8(0.6, 1.0) |
| Grade |  |  |  |  |  |  |  |  |  |  |  |  |  |  |  |  |  |  |  |
| I | 168 | 24(19.8) | 15.2 | (Reference) | 19(18.4) | 12.0 | (Reference) | 3(9.1) | 1.9 | (Reference) | 21(14.1) | 13.3 | (Reference) | 19(22.6) | 12.0 | (Reference) | 36(17.6) | 22.8 | (Reference) |
| II | 596 | 83(68.6) | 14.0 | 0.9(0.6, 1.4) | 77(74.8) | 13.0 | 1.1(0.7, 1.7) | 27(81.8) | 4.6 | 2.4(0.7, 7.8) | 116(77.9) | 19.5 | 1.5(1, 2.3) | 56(66.7) | 9.4 | 0.8(0.5, 1.3) | 138(67.6) | 23.3 | 1.0(0.7, 1.4) |
| III | 90 | 13(10.7) | 14.6 | 1.0(0.5, 1.8) | 7(6.8) | 7.9 | 0.7(0.3, 1.5) | 2(6.1) | 2.2 | 1.2(0.2, 7.0) | 12(8.1) | 13.5 | 1.0(0.5, 2.0) | 8(9.5) | 9.0 | 0.7(0.3, 1.6) | 30(14.7) | 33.7 | 1.5(1.0, 2.2) |
| IV | 7 | 1(0.8) | 14.3 | 0.9(0.1, 6.0) | 0(0) | - | - | 1(3.0) | 14.3 | 7.5(0.9, 63.5) | 0(0) | - | - | 1(1.2) | 14.3 | 1.2(0.2, 7.7) | 0(0) | - | - |
| Stage |  |  |  |  |  |  |  |  |  |  |  |  |  |  |  |  |  |  |  |
| I | 168 | 25(16.9) | 15.0 | (Reference) | 18(14.8) | 10.8 | (Reference) | 6(14) | 3.6 | (Reference) | 23(13.5) | 13.8 | (Reference) | 17(16.5) | 10.2 | (Reference) | 34(14.4) | 20.4 | (Reference) |
| II | 281 | 51(34.5) | 18.4 | 1.2(0.8, 1.9) | 31(25.4) | 11.2 | 1.0(0.6, 1.8) | 13(30.2) | 4.7 | 1.3(0.5, 3.4) | 57(33.3) | 20.6 | 1.5(1.0, 2.3) | 42(40.8) | 15.2 | 1.5(0.9, 2.5) | 69(29.2) | 25.0 | 1.2(0.9, 1.8) |
| III | 285 | 29(19.6) | 10.4 | 0.7(0.4, 1.1) | 39(32.0) | 13.9 | 1.3(0.8, 2.2) | 11(25.6) | 3.9 | 1.1(0.4, 2.9) | 54(31.6) | 19.3 | 1.4(0.9, 2.2) | 16(15.5) | 5.7 | 0.6(0.3, 1.1) | 79(33.5) | 28.2 | 1.4(1.0, 2.0) |
| IV | 267 | 43(29.1) | 16.2 | 1.1(0.7, 1.7) | 34(27.9) | 12.8 | 1.2(0.7, 2.0) | 13(30.2) | 4.9 | 1.4(0.5, 3.5) | 37(21.6) | 14.0 | 1.0(0.6, 1.6) | 28(27.2) | 10.6 | 1.0(0.6, 1.8) | 54(22.9) | 20.4 | 1.0(0.7, 1.5) |
| Healthcare factors | Total |  |  |  |  |  |  |  |  |  |  |  |  |  |  |  |  |  |  |
| Type hospital admission |  |  |  |  |  |  |  |  |  |  |  |  |  |  |  |  |  |  |  |
| Emergency | 183 | 25(22.3) | 14.0 | (Reference) | 19(19.2) | 10.6 | (Reference) | 10(33.3) | 5.6 | (Reference) | 34(23.3) | 19.0 | (Reference) | 14(16.1) | 7.8 | (Reference) | 30(14.7) | 16.8 | (Reference) |
| Planned | 693 | 87(77.7) | 12.7 | 0.9(0.6, 1.4) | 80(80.8) | 11.7 | 1.1(0.7, 1.8) | 20(66.7) | 2.9 | 0.5(0.2, 1.1) | 112(76.7) | 16.4 | 0.9(0.6, 1.2) | 73(83.9) | 10.7 | 1.4(0.8, 2.4) | 174(85.3) | 25.5 | 1.5(1.1, 2.2) |
| Surgery |  |  |  |  |  |  |  |  |  |  |  |  |  |  |  |  |  |  |  |
| Yes | 879 | 113(73.9) | 13.0 | (Reference) | 100(80.6) | 11.5 | (Reference) | 30(63.8) | 3.5 | (Reference) | 147(80.8) | 17.0 | (Reference) | 88(84.6) | 10.2 | (Reference) | 206(83.1) | 23.8 | (Reference) |
| No | 175 | 40(26.1) | 23.3 | 1.8(1.3, 2.5) | 24(19.4) | 14.0 | 1.2(0.8, 1.8) | 17(36.2) | 9.9 | 2.8(1.6, 5.0) | 35(19.2) | 20.3 | 1.2(0.9, 1.7) | 16(15.4) | 9.3 | 0.9(0.6, 1.5) | 42(16.9) | 24.4 | 1.0(0.8, 1.4) |
| Type of Surgery |  |  |  |  |  |  |  |  |  |  |  |  |  |  |  |  |  |  |  |
| Major | 816 | 108(95.6) | 13.4 | (Reference) | 93(93.9) | 11.6 | (Reference) | 28(96.6) | 3.5 | (Reference) | 140(96.6) | 17.4 | (Reference) | 79(90.8) | 9.8 | (Reference) | 193(96.5) | 24.1 | (Reference) |
| Minor | 43 | 5(4.4) | 11.6 | 0.9(0.4, 2.0) | 6(6.1) | 14.0 | 1.2(0.6, 2.6) | 1(3.4) | 2.3 | 0.7(0.1, 4.8) | 5(3.4) | 11.6 | 0.7(0.3, 1.5) | 8(9.2) | 18.6 | 1.9(1.0, 3.7) | 7(3.5) | 16.3 | 0.7(0.3, 1.3) |
| Time to surgery in months |  |  |  |  |  |  |  |  |  |  |  |  |  |  |  |  |  |  |  |
| Emergency 0 days | 171 | 16(14.3) | 9.5 | (Reference) | 15(15.2) | 8.9 | (Reference) | 9(30.0) | 5.4 | (Reference) | 21(14.4) | 12.5 | (Reference) | 11(12.6) | 6.5 | (Reference) | 32(15.6) | 19.0 | (Reference) |
| 1 to <14 days | 115 | 16(14.3) | 14.3 | 1.5(0.8, 2.9) | 10(10.1) | 8.9 | 1.0(0.5, 2.1) | 5(16.7) | 4.5 | 0.8(0.3, 2.4) | 19(13.0) | 17.0 | 1.4(0.8, 2.4) | 11(12.6) | 9.8 | 1.5(0.7, 3.3) | 18(8.8) | 16.1 | 0.8(0.5, 1.4) |
| 14 to 30 days | 124 | 13(11.6) | 11.0 | 1.2(0.6, 2.3) | 8(8.1) | 6.8 | 0.8(0.3, 1.7) | 4(13.3) | 3.4 | 0.6(0.2, 2.0) | 24(16.4) | 20.3 | 1.6(1.0, 2.8) | 11(12.6) | 9.3 | 1.4(0.6, 3.2) | 29(14.1) | 24.6 | 1.3(0.8, 2.0) |
| 31 to 59 days | 188 | 26(23.2) | 13.8 | 1.5(0.8, 2.6) | 28(28.3) | 14.9 | 1.7(0.9, 3.0) | 7(23.3) | 3.7 | 0.7(0.3, 1.8) | 32(21.9) | 17.0 | 1.4(0.8, 2.3) | 28(32.2) | 14.9 | 2.3(1.2, 4.4) | 56(27.3) | 29.8 | 1.6(1.1, 2.3) |
| 60 and more days | 280 | 41(36.6) | 14.9 | 1.6(0.9, 2.7) | 38(38.4) | 13.8 | 1.6(0.9, 2.8) | 5(16.7) | 1.8 | 0.3(0.1, 1.0) | 50(34.2) | 18.1 | 1.5(0.9, 2.3) | 26(29.9) | 9.4 | 1.5(0.8, 2.9) | 70(34.1) | 25.5 | 1.3(0.9, 1.9) |

P: Prevalence; Comorbidities: II Congestive heart failure; III Peripheral vascular disease; V Dementia; VI Chronic pulmonary disease; VII Rheumatic disease; IX Diabetes mellitus

### Distribution and frequency of comorbidities by tumor characteristics

The pattern of comorbidities by region was similar, except for two comorbidities (rheumatologic disease and congestive heart failure), where Granada presented a higher prevalence than Girona. The pattern of comorbidities by sex shows a high prevalence of COPD among male colorectal cancer patients (79%), while almost 60% of patients with dementia or rheumatologic disease were female. There was a frequency gradient of comorbidities by age, with dementia (75%), congestive heart failure (64%), and renal disease (46%) as the most common comorbidities among the elderly. Patients’ performance status varied among comorbidities as well. Ninety-two percent of liver disease patients and 80% of diabetes patients had ECOG performance score 0 or 1, in contrast to only 53% of dementia and 30% of congestive heart failure patients. There was strong evidence supporting a significant trend of comorbidity prevalence across the levels of performance status for the five most common comorbidities plus dementia. Furthermore, COPD, diabetes, and dementia were more frequently associated with smoking (current and previous): 68%, 53%, and 36%, respectively. Adjusted PRs (APRs) comparing current smoker vs. never smoker in COPD, diabetes, and dementia were 3.1 (95% CI: 1.9-5.0), 1.3 (95% CI: 0.8-2.0), and 1.8 (95% CI: 0.6-5.2), respectively. Overweight and obesity were more prevalent among patients with congestive heart failure (81%), peripheral vascular disease (76%), and diabetes (77%). The respective comorbidity APRs comparing a BMI ≥30 kg/m^2^ vs. <25 kg/m^2^ were 2.1 (95% CI: 1.2-3.6) for congestive heart failure, 1.7 (95% CI: 1.0-2.7) for peripheral vascular disease, and 1.7 (95% CI: 1.2-2.4) for diabetes. However, patients with dementia showed the highest prevalence of underweight and normal weight (body mass index <25 kg/m^2^) patients (41%) (Tables 2 and S2).

### Distribution and frequency of comorbidities by tumor’s characteristics

Among patients with dementia the most common anatomical site was the right side of the colon (44%). However, among patients with rheumatic disease the most common anatomical site was the rectum (38%). Regarding the grade of differentiation, the most common grade for all the different comorbidities was grade two (moderately differentiated). However, patients with diabetes had the highest proportion of grade three (30%) and an APR of 1.4 (95% CI: 0.9-2.0) comparing grades three-four vs. one. Overall, all comorbidities had approximately 55% of cancer cases diagnosed at stages III or IV. Patients with COPD showed the lowest frequency of stage IV (22%). CRC patients with dementia had a 30% higher prevalence of advanced cancer diagnosis i.e. APR 1.3; 95% CI: 0.5-3.2 comparing stage IV vs I (Tables 2 and S2).

### Distribution and frequency of comorbidities by healthcare characteristics

Patients with dementia showed the highest prevalence of emergency hospital admission after CRC diagnosis (33%) with an APR comparing planned vs. emergency admission of 1.6 (95% CI: 1.1-2.2). Despite the emergency admission, dementia was the comorbidity with the highest prevalence of patients who were not receiving surgery as treatment (64%) with an APR of 2.1 (95% CI: 1.2-3.8). Note that patients with dementia also showed the second highest prevalence of stage IV, with 30% of the cases. However, patients with rheumatologic disease showed the highest prevalence of major surgery (91%) and also the highest APR for minor surgery (2.0; 95% CI: 1.0-3.7). Major surgery was the most common type of surgery among all CRC patients, with at least 90% for all comorbidities. The pattern of time to surgery by comorbidities showed considerable variability. Overall, among the majority of comorbidities, one-third of CRC patients were offered surgery 60 or more days after the cancer diagnosis. However, dementia patients showed a different pattern: 30% had emergency surgery the same day as hospital admission (time to surgery of zero days). CRC with congestive heart failure showed the highest APR (1.7; 95% CI: 1.0-2.9) comparing surgery more than 60 days vs. emergency surgery (zero days) (Tables 2 and S2).

Table 4 shows the prevalence ratios for the presence of multimorbidity versus the absence of comorbidities by patients, tumor, and healthcare factors. Overall, a higher prevalence of multimorbidity was associated with being aged ≥75 years, obese, male, or current smoker (Figure 3).

**Table 4.**
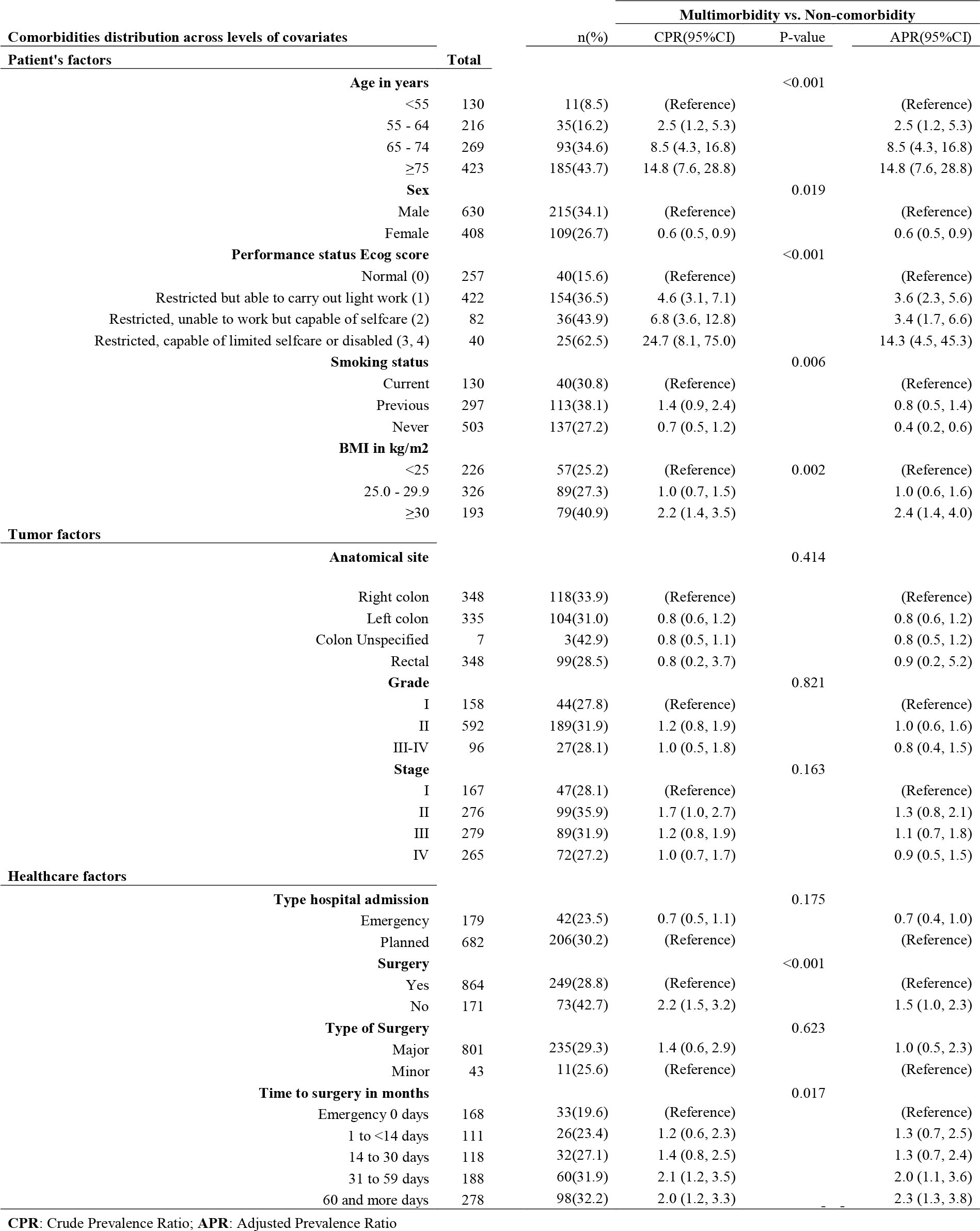
Multimorbidity prevalence ratios by patient, tumor and healthcare characteristics among all incident colorectal cancer patients in Granada and Girona, 2011, n = 1,061.

**Figure 3.**
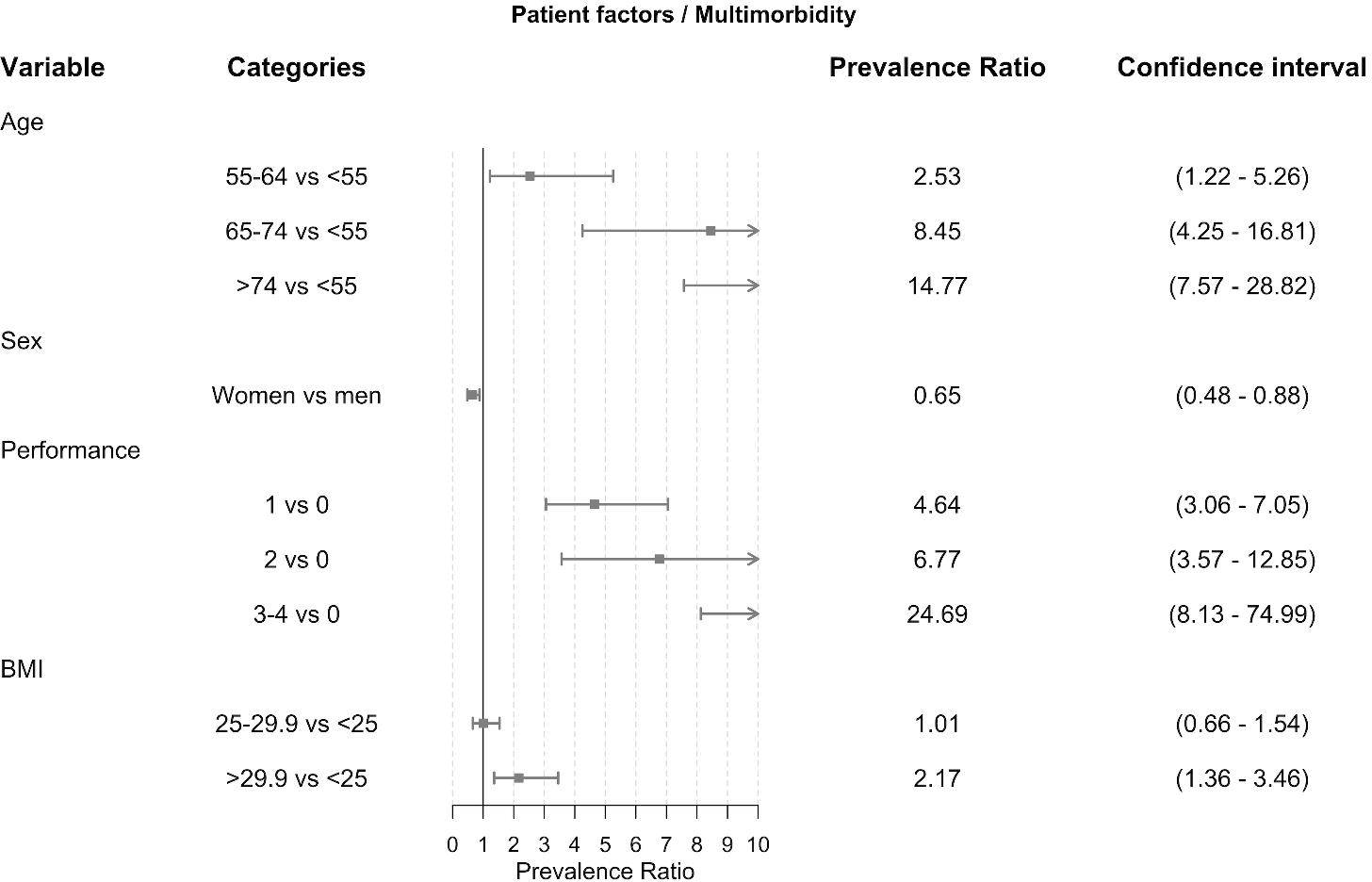
Forest plot: Multimorbidity prevalence ratios by patients’ age, sex, performance status and BMI among all incident colorectal cancer patients in Granada and Girona, 2011, n = 1,061.

Likewise, being offered surgery more than 60 days after cancer diagnosis and not receiving surgery were associated with a higher risk of multimorbidity. It is important to highlight that 37% of patients having emergency surgery had multimorbidity and were aged ≥75 years. Furthermore, 30% of emergency surgery was performed in older (≥75 years) advanced stage (III/IV) CRC patients affected by dementia. There was moderated evidence supporting that patients with multimorbidity versus non-comorbidity had a 50% higher prevalence of not receiving surgery (APR 1.5; 95% CI: 1.0-2.3). However, we found strong evidence of receiving surgery more than 60 days after cancer diagnosis in CRC patients with multimorbidity compared to patients free of comorbidities. Patients affected by multimorbidity had 2.3 times higher prevalence of receiving late surgery compared to emergency surgery (0 days) (APR: 2.3; 95% CI: 1.3-3.8) (Table 4).

Furthermore, the complete visualization of CoMCoR study results is provided at the following link http://watzilei.com/shiny/CoMCoR/.

## Discussion

Overall, comorbidity is commonly recognized as being associated with cancer outcomes and survival [23]. However, there is an international sparsity of population-based epidemiological studies describing the prevalence of comorbidities and multimorbidity among cancer patients [24]. CoMCoR study fills this gap, providing translating knowledge into clinical practice regarding the pattern of the prevalence of comorbidities and multimorbidity among CRC patients in Spain. The pattern is mainly characterized by a higher prevalence of diabetes, advanced cancer stage, and late surgery or no surgical treatment in older patients with dementia.

To the best of our knowledge, the CoMCoR study presented here is the first to identify the most prevalent comorbidities and multimorbidity among CRC patients in Spain, and characterize a particular pattern in the distribution and frequency of comorbidities and multimorbidity. While clinical studies are representative of only a selected part of the population, CoMCoR is a population-based observational study using cancer registration and hospital medical records that translates its results into clinical practice based on real-world data.

Regarding the prevalence of comorbidities, we found that diabetes is the most prevalent comorbidity among CRC patients (24%). Among non-cancer populations, the prevalence of any type of diabetes in adults in Spain has been reported to range between 6 and 11% [25]. However, there is a scarcity of literature reporting the prevalence of diabetes among CRC patients [24]. Our findings were similar to those previously reported in a Taiwanese cohort of 1,197 CRC patients where 24% had either a reported history of diabetes or were currently taking one or more diabetes-controlling medications [26]. Some evidence shows that diabetes is associated with higher incidence of CRC and shorter CRC survival [27]. Thus, we argue that public health programs targeting cancer prevention strategies among diabetic patients might have a positive impact on CRC outcomes in Spain.

Furthermore, we found a high prevalence of advanced stage cancer diagnosis (stage III/IV) among all CRC patients, which was even higher in older CRC patients affected by dementia compared to the prevalence we would have expected had they been offered the screening. Recently in Denmark it has been shown that CRC patients that had offered the screening lower prevalence of advance cancer stage at diagnosis [28]. . We argue that this may be due to low utilization of CRC screening in Spain. In 2011, CRC screening programs were implemented in only nine Spanish regions, with just partial coverage [29]. While all populations would benefit from the systematic use of screening, socioeconomically disadvantaged groups, such as patients with dementia, may especially benefit from a targeted CRC screening [30]. We acknowledge that our study design does not allow for recommendations about targeted CRC screening. However, further research might explore the impact of the screening among colorectal cancer patients in Spain.

Comorbid medical diseases are highly prevalent among elderly. Overall, over 60% of all cases of cancer are diagnosed after age 65 years, with 67% of cancer deaths occurring in this age group [31]. We found a high prevalence of older patients not receiving surgical treatment, but it was even higher for older patients with stage III/IV CRC and dementia. There are many reasons why cancer occurs more frequently in older persons. The elderly have less resistance and longer exposure to carcinogens, a decline in immune system functioning, an alteration in anti-tumor defenses, decreased DNA repair, defects in tumor-suppressor genes, and differences in biological behavior, including angiogenesis. These factors contribute to the elderly population often being affected by comorbidities which affect cancer diagnosis, treatment, and survival [32]. The high prevalence we found of older CRC patients not receiving surgical treatment in stages III and IV in addition to a higher prevalence of contraindications it partially might reflect the low uptake and partial coverage of CRC screening and preventive strategies in Spain.

Regarding multimorbidity, we found that it is associated with late surgery (≥60 days after cancer diagnosis) and emergency surgery offered the same day of an emergency hospital admission. Recently published evidence has shown that CRC diagnosed after a hospital emergency room admission were more likely associated with older and more socioeconomically deprived individuals [33]. Although disease stage at the time of diagnosis of CRC is a crucial determinant of patient outcome, comorbidity increases the complexity of cancer management and affects survival duration. Cancer control and treatment research questions should address multimorbidity, particularly in the elderly [34]. Regarding the evidence examining time from cancer diagnosis to surgical treatment there is no conclusive evidence supporting an optimal window of time. However, a study from the American College of Surgeons has found that patients who had a cancer operation at precisely eight weeks (56 days) after the end of combined chemoradiotherapy had the best overall survival and successful removal of their residual tumors [35]. Other study found that CRC patients waiting longer than 12 weeks (84 days) to receive surgery had increased all-cause mortality compared with patients receiving surgery within four weeks (28 days) [36]. In a study of patients receiving elective surgery for colonic resection following diagnosis with CRC in Ontario, it was found that factors influencing receipt of treatment after 42 days from diagnosis included older age and comorbidity [37].

Emergency surgery was defined as surgery offered the same day of an emergency hospital admission. Thus, we were assuming implicitly that CRC was diagnosed as a consequence of an emergency surgical intervention. However, we do not have empirical data to support our assumption. On the other hand, 30% of emergency surgery was performed among older advanced-stage CRC patients with dementia. It has been shown that CRC diagnosed after a hospital emergency admission is more likely associated with older and more deprived individuals [38, 39]. Recently, a study showed that 18% of CRC cases that were diagnosed as emergency cases had “red flag” symptoms, indicating the disease could have been identified earlier [33]. The promotion of CRC symptom awareness among the elderly and the caregivers of older patients affected from dementia might help them to early identify these symptoms and visit their general practitioner, who must refer them through the normal pathways to specialist evaluation [33].

There have been attempts to reanalyze the different comorbidity scores and their weighting algorithms, which show that some diseases should have a higher weight (including dementia), and others a lower weight (including peptic ulcers). Different approaches to measuring comorbidity specifically in cancer patients include focusing on single comorbid conditions in isolation, or weighted indices such as the Charlson comorbidity index [40], the Adult Comorbidity Evaluation – 27 index (ACE-27) [41], or the Elixhauser index [42]. However, to date, there is no agreed gold standard method upon which to measure comorbidity in the cancer patient population [43]. We used the Royal College of Surgeons system, which is a clinical score used to evaluate the risk of death during surgery. The score applies an equal weight system to 12 different comorbidities categorized into 0, 1, 2 or more comorbidities, making it easy-to-use, since all comorbidities are considered equally important [20].

We assumed that missing data were completely at random and performed a complete case analysis, which might introduce bias if the data were actually missing at random. However, our CoMCoR study was merely descriptive, and the percentage of missing data for the main outcome (comorbidities) was only 2%. Also, we would like to acknowledge the limited scope of the analysis in terms of time and space, with only one calendar year of CRC incident cases and two population-based cancer registries, thus limiting the external validity of our findings and supporting the need of more studies.

In summary, the CoMCoR study has identified existing patterns in the distribution and frequency of comorbidities and multimorbidity for CRC patients in Spain by patient, tumor and health care factors. The high prevalence of CRC diagnosed at stage III/IV among elderly patients and patients with dementia and the high prevalence of older patients not receiving surgical treatment are significant findings that require immediate policy actions. Results from the CoMCoR study may help to foster CRC screening and preventive strategy policies in Spain and other countries.

## Acknowledgments

We thank Minicozzi Pamela and Sant Milena for the development of the protocol and data recollection tools for the European High-Resolution studies.

## Supporting information

**Supplementary Table S1.** Risk factors associated with the top-five comorbidities adjusted by sex and age among all incident colorectal cancer patients by patient characteristics during 2011 in Granada and Girona, n = 1,061

**Supplementary Table S2.** Distribution and frequency of comorbidities by patient’s characteristics among all incident colorectal cancer patients in Granada and Girona, 2011, n = 1,061

**Supplementary Table S3.** Distribution and frequency of comorbidities by tumor characteristics among all incident colorectal cancer patients in Granada and Girona, 2011, n = 1,061

**Supplementary Table S4**. Distribution and frequency of comorbidities by healthcare characteristics among all incident colorectal cancer patients in Granada and Girona, 2011, n = 1,061

